# Different pattern of pre-existing SARS-COV-2 specific T cell immunity in SARS-recovered and uninfected individuals

**DOI:** 10.1101/2020.05.26.115832

**Authors:** Nina Le Bert, Anthony T Tan, Kamini Kunasegaran, Christine Y L Tham, Morteza Hafezi, Adeline Chia, Melissa Chng, Meiyin Lin, Nicole Tan, Martin Linster, Wan Ni Chia, Mark I-Cheng Chen, Lin-Fa Wang, Eng Eong Ooi, Shirin Kalimuddin, Paul Anantharajal Tambyah, Jenny Guek-Hong Low, Yee-Joo Tan, Antonio Bertoletti

## Abstract

Memory T cells induced by previous infections can influence the course of new viral infections. Little is known about the pattern of SARS-CoV-2 specific pre-existing memory T cells in human. Here, we first studied T cell responses to structural (nucleocapsid protein, NP) and non-structural (NSP-7 and NSP13 of ORF1) regions of SARS-CoV-2 in convalescent from COVID-19 (n=24). In all of them we demonstrated the presence of CD4 and CD8 T cells recognizing multiple regions of the NP protein. We then show that SARS-recovered patients (n=23), 17 years after the 2003 outbreak, still possess long-lasting memory T cells reactive to SARS-NP, which displayed robust cross-reactivity to SARS-CoV-2 NP. Surprisingly, we observed a differential pattern of SARS-CoV-2 specific T cell immunodominance in individuals with no history of SARS, COVID-19 or contact with SARS/COVID-19 patients (n=18). Half of them (9/18) possess T cells targeting the ORF-1 coded proteins NSP7 and 13, which were rarely detected in COVID-19- and SARS-recovered patients. Epitope characterization of NSP7-specific T cells showed recognition of protein fragments with low homology to “common cold” human coronaviruses but conserved among animal betacoranaviruses.

Thus, infection with betacoronaviruses induces strong and long-lasting T cell immunity to the structural protein NP. Understanding how pre-existing ORF-1-specific T cells present in the general population impact susceptibility and pathogenesis of SARS-CoV-2 infection is of paramount importance for the management of the current COVID-19 pandemic.

## Main Text

Severe acute respiratory syndrome coronavirus-2 (SARS-CoV-2) is the cause of the coronavirus disease 2019 (COVID-19)^1^. This disease has spread pandemically placing lives and economies of the world under severe stress. SARS-CoV-2 infection is characterized by a broad spectrum of clinical syndromes, ranging from mild influenza-like symptoms to severe pneumonia and acute respiratory distress syndrome^2^.

It is common to observe in human the ability of a single virus to cause different pathological manifestations. This is often due to multiple contributory factors including the quantity of viral inoculum, the genetic background of patients and the presence of concomitant pathological conditions. Moreover, an established adaptive immunity towards closely related or completely different viruses can increase protection^3^ or enhance disease severity^4^.

SARS-CoV-2 belongs to *Coronaviridae,* a family of large RNA viruses infecting many animal species. Six other coronaviruses are known to infect human. Four of them are endemically transmitted^5^ and cause common cold (OC43, HKU1, 229E and NL63), while SARS-CoV (defined from now as SARS-CoV-1) and MERS-CoV have caused limited epidemics of severe pneumonia^6^. All of them trigger antibody and T cell responses in infected patients: however, antibody levels appear to wane relatively quicker than T cells. In SARS recovered patients, SARS-CoV-specific antibodies dropped below detection limit within 2 to 3 years^7^, while SARS-CoV-specific memory T cells can be detected even at 11 years after infection^8^. Since the sequences of selected structural and non-structural proteins are highly conserved among different coronaviruses (i.e. NSP7 and NSP13 are 100% and 99% identical, respectively, between SARS-CoV-2, SARS-CoV-1 and the bat-SL-CoVZXC21^9^), we studied whether crossreactive SARS-CoV-2-specific T cells are present in individuals who resolved from SARS-CoV-1 or SARS-CoV-2 infection. We also studied these T cells in individuals with no history of SARS or COVID-19 and who were also not in contact with SARS-CoV-2 infected cases. Collectively these individuals are hereon referred to as SARS-CoV-1/2 unexposed.

SARS-CoV-2-specific T cells have just started to be characterized in COVID-19 patients^10,11^ and their potential protective role has been inferred from studies in SARS^12^ and MERS^13^ patients. To study SARS-CoV-2 specific T cells associated with viral clearance, we collected peripheral blood of 24 individuals who recovered from mild to severe COVID-19 (demographic, clinical and virological information are summarized in **Extended Data Table 1**) and studied the T cell response against selected structural (nucleocapsid protein-NP) and non-structural proteins (NSP7 and NSP13 of ORF1) of the large SARS-CoV-2 proteome (**Figure 1A**). We selected nucleocapsid protein as it is one of the more abundant structural proteins produced and has large homology between different betacoranaviruses (**Extended Data Fig. 1**)^14^.

**Fig 1:**
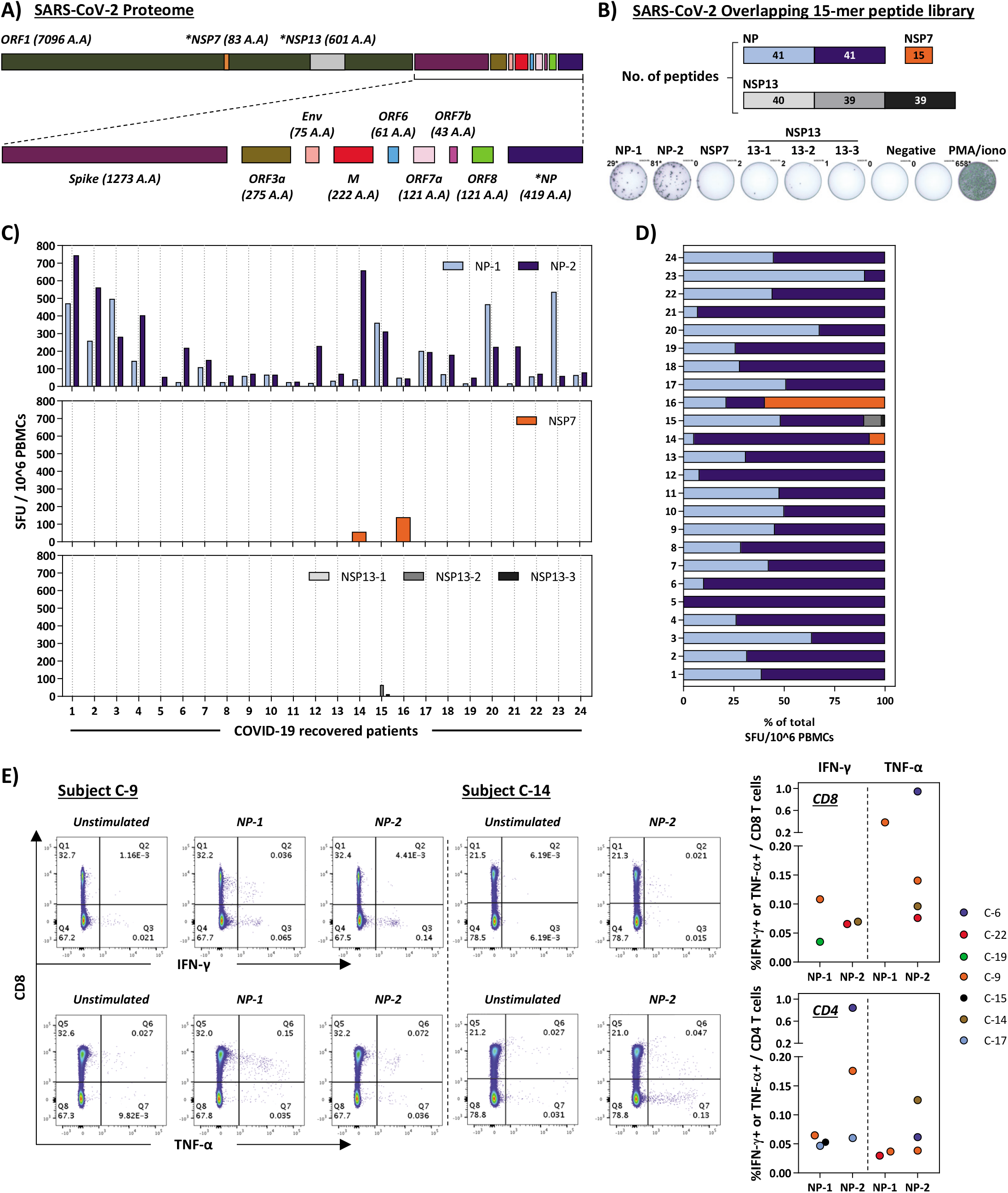
SARS-CoV-2-specific T cells in recovered COVID-19 patients. (A) SARS-CoV-2 proteome organization; analyzed proteins are marked by *. (B) For detection of SARS-CoV-2-specific T cells by IFN-γ ELISpot, 15-mer peptides overlapping by 10 amino acids covering nucleocapsid protein (NP) and the non-structural proteins (NSP) 7 and 13 were synthesized and split into 5 pools of about 40 peptides covering NP (NP-1, NP-2) and NSP13 (NSP13-1, NSP13-2, NSP13-3) and one pool of 15 peptides covering NSP7. (C) PBMC of 24 recovered COVID-19 patients were stimulated with the peptide pools. Bar graphs show frequency of spot forming units (SFU) of IFN-γ secreting cells. (C) Composition of the SARS-CoV-2-specific T cell repertoire is shown as percentage of SARS-CoV-2-specific T cells reacting to NP (NP-1 = light blue; NP2 = dark blue), NSP7 (orange) and NSP13 (grey) for the individual recovered COVID-19 patients tested. (D) PBMC were stimulated with the peptide pools covering NP (NP-1, NP-1) for 5h and analyzed by intracellular cytokine staining. Dot plots show examples of patients with CD4 and/or CD8 T cells producing IFN-γ and/or TNF-α in response to stimulation with NP-1 and/or NP2 peptides. The graphs summarize the percentage of SARS-CoV-2 NP-peptide-reactive CD4 and CD8 T cells in 7 individuals.

NSP7 and NSP13 were selected for their complete homology between SARS-CoV-1, SARS-CoV-2 and other animal coronaviruses belonging to the betacoranavirus genus (**Extended Data Fig. 2**)^9^, and because they are representative of the ORF1a/b polyprotein encoding the replicase-transcriptase complex^15^. This polyprotein is the first to be translated upon coronavirus infection. We synthesized 216 15-mer peptides overlapping by 10 amino acids (aa) covering the whole length of NSP7 (83aa), NSP13 (601aa) and NP (422aa) that were organized in 5 pools of approximately 40 peptides each (NP-1, NP-2, NSP13-1, NSP13-2, NSP13-3) and in a single pool of 15 peptides spanning NSP7 (**Figure 1B**). The unbiased method with overlapping peptides was utilized instead of peptide selection by bioinformatic approaches, since the performance of such algorithms in ethnically-diverse Asians is often sub-optimal^16^.

Peripheral blood mononuclear cells (PBMC) of 24 recovered COVID-19 patients were stimulated for 18h with the different peptide pools and virus-specific T cell responses were analyzed by IFN-γ ELISpot assay. In all tested individuals (24/24) we detected IFN-γ spots following stimulation with the pools of synthetic peptides covering NP (**Figure 1C/D**). In nearly all individuals NP-specific responses could be identified for multiple regions of the protein: 23/24 for region 1-205aa (NP-1) and 24/24 for 206-422aa (NP-2). In sharp contrast, responses to NSP7 and NSP13 peptide pools were detected at low levels only in 3 out of 24 COVID-19 convalescents tested. Direct *ex vivo* intracellular cytokine staining (ICS) was performed to confirm and define the NP-specific IFN-γ ELISpot response. Due to the low frequency, NP-specific T cells were more difficult to visualize by ICS than by ELISpot, but a clear population of CD4 and/or CD8 T cells producing IFN-γ and/or TNF-α were detectable in 7 out of 9 tested subjects (**Figure 1E**). To confirm and further delineate the multispecificity of the NP-specific T cell response detected *ex vivo* in COVID-19 recovered patients, we defined in nine individuals, the distinctive sections of NP targeted by T cells. We organized the 82 overlapping peptides covering the entire NP into small peptide pools (7-8 peptides) that were used to stimulate PBMC either directly *ex vivo* or after an *in vitro* expansion protocol previously used in HBV^17^ or SARS recovered subjects^18^. A schematic representation of the peptide pools is shown in **Figure 2A**. We found that 8 out of 9 COVID-19 recovered patients possess T cells that recognize multiple regions of NP of SARS-CoV-2 (**Figure 2A**). Importantly, we then defined single peptides that were able to activate T cells in 7 patients. Utilizing a peptide matrix strategy^18^, we first deconvolute individual peptides responsible for the detected T cell response by IFN-γ ELISpot. Subsequently, we confirmed the identified single peptide by testing, with ICS, its ability to activate CD4 or CD8 T cells (**Figure 2B**). **Figure 2B** summarizes the different T cell epitopes defined by both ELISpot and ICS, in 7 COVID-19 recovered individuals. Remarkably, we observed that COVID-19 convalescents developed T cells specific to regions that were also targeted by T cells of SARS recovered subjects. For example, the NP region 101-120 which is a described CD4 T cell epitope in SARS-CoV-1 exposed individuals^8,18^, also stimulated CD4 T cells of two COVID-19 recovered donors. Similarly, the NP region 321-340 contains epitopes triggering CD4 and CD8 T cells in both COVID-19 and SARS recovered patients^18^. The demonstration that COVID-19 and SARS recovered patients can mount T cell responses against shared viral determinants implicates that individuals with SARS-CoV-1 infection can induce T cells able to cross-react against SARS-CoV-2.

**Fig 2:**
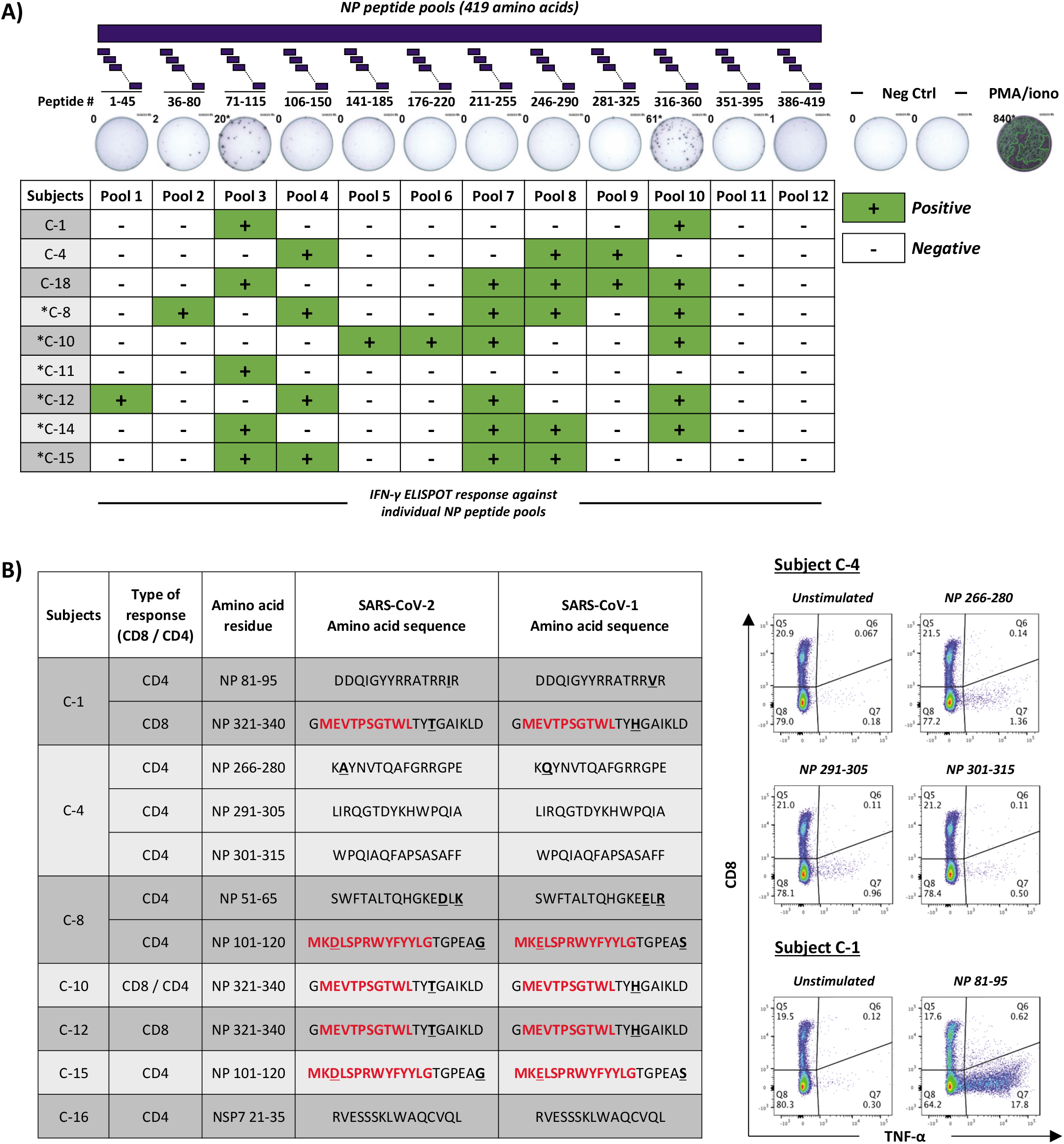
SARS-CoV-2-specific T cells in convalescent COVID-19 patients are targeting multiple regions of nucleocapsid protein. (A) PBMC of 9 COVID-19 recovered individuals were stimulated with 12 different pools of 7-8 NP-peptides to delineate the region to which their T cells react. The table shows IFN-γ ELISpot response against the individual NP peptide pools. (B) Following in vitro T cell expansion, a peptide pool matrix strategy was applied in 7 individuals. The distinct peptide epitopes to which the T cells react were identified by IFN-γ ELISpot and confirmed by ICS (representative dot-plots are shown). Previously described T cell epitopes for SARS-CoV-1 are highlighted in red; non-conserved amino acid residues between SARS-CoV-1 and −2 are bold and underlined.

For the management of the current pandemic and for vaccine development against SARS-CoV-2, it is important to understand if acquired immunity will be long-lasting. Therefore, we tested if individuals who recovered from SARS 17 years ago still harbor memory T cells against SARS-CoV-1. Hence, their PBMC (n=15) were stimulated directly *ex vivo* with peptide pools covering SARS-CoV-1 NP (NP-1 and NP-2), NSP7 and NSP13 (**Figure 3A**). This revealed that 17 years after infection, those individuals still possess virus-specific memory T cells, and similar to COVID-19 recovered patients, we detected T cells reacting almost exclusively to NP and not to the NSPs (**Figure 3B/C**). Subsequently, we tested if the NP-specific T cells detected in SARS recovered patients could cross-react with SARS-CoV-2 NP peptides (aa identity = 94%). Indeed, although at lower frequency, T cells in all 23 individuals tested reacted to SARS-CoV-2 NP (**Figure 3D, 4A**). In order to test whether these T cells could expand after encounter with SARS-CoV-2 NP, their PBMC were stimulated *in vitro* with the whole battery of NP, NSP7 and NSP-13 peptides and the quantity of T cells responding to SARS-CoV-2 NP, NSP7 and NSP13 was analyzed after 10 days of cell culture. A clear and robust expansion of NP-specific T cells was detected in 7 out of 8 individuals tested (**Figure 3E**). Importantly, and in sharp contrast to the T cell response to NP peptides, we could not detect any T cells reacting to the peptide pools covering NSP13 and only 1 out of 8 reacted to NSP7, despite *in vitro* expansion.

**Fig 3:**
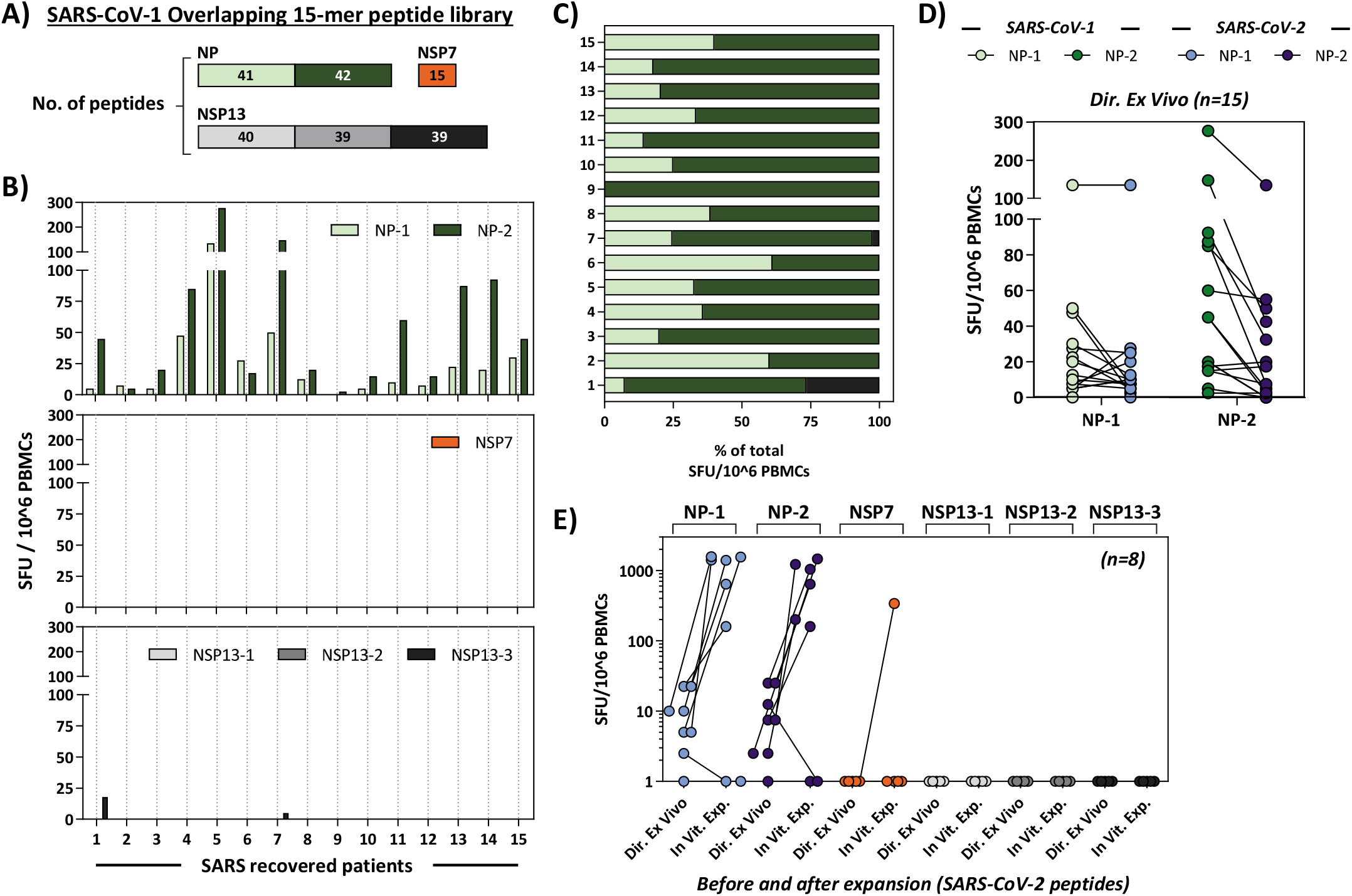
SARS-CoV-2 cross-reactive memory T cells are present in SARS-recovered patients. (A) PBMC isolated from 15 individuals who recovered from SARS 17 years ago were stimulated with SARS-CoV-1 NP, NSP7 and NSP13 peptide pools. (B) Bar graphs show spot forming units (SFU) of IFN-γ secreting cells per 1 million PBMC following overnight stimulation with the indicated peptide pools. (C) Composition of the SARS-CoV-1-specific T cells in individual recovered SARS patients. The percentage of SARS-CoV-1-specific T cells against NP (NP-1 = light green; NP2 = dark green), NSP7 (orange) and NSP13 (grey) in each patient is shown. (D) PBMC of 15 SARS-recovered individuals were stimulated in parallel with peptide pools covering NP of SARS-CoV-1 and of SARS-CoV-2 and their frequency is shown. (E) PBMC of 8 SARS-recovered individuals were stimulated with all peptides covering SARS-CoV-2 NP, NSP7 and NSP13 to expand peptide cross-reactive T cells. The graph shows the number of T cells reactive to the peptide pools indicated directly ex-vivo and after specific T cell expansion.

Thus, SARS-CoV-2 NP-specific cross-reactive T cells are part of the T cell repertoire of individuals with a history of SARS-CoV-1 infection and are able to robustly expand after encounter with SARS-CoV-2 NP peptides. These findings demonstrate that virus-specific memory T cells induced by betacoronanvirus infection are long-lasting, which supports the notion that COVID-19 patients would develop long-term T cell immunity. Furthermore, our findings also raise the intriguing possibility that infection with related viruses can also protect from or modify the pathology caused by SARS-CoV-2 infection.

To explore this possibility, we tested NP and NSP7/13-specific T cell responses in 18 SARS-CoV-1/2 unexposed donors. The blood samples were collected either before July 2019 or were serologically negative for both SARS-CoV-2 neutralizing antibodies and SARS-CoV-2 NP antibodies^19^. Different coronaviruses known to cause common cold in humans like OC43, HKU1, NL63 and 229E present different degrees of amino acid homology with SARS-CoV-2 (**Extended Data Fig. 1, 2**) and recent data demonstrated the presence of SARS-CoV-2 cross-reactive CD4 T cells (mainly specific for Spike) in SARS-CoV-2 unexposed donors^11^. Remarkably, we detected NP-specific T cells in some of our SARS-CoV-1/2 unexposed individuals. The pattern of T cell reactivity, however, was different compared to COVID-19 and SARS recovered. T cells from SARS-CoV-1/2 unexposed were directed against a single peptide pool: i.e. none of the 18 donors responded to the NP-2 peptide pool (**Figure 4A**). Moreover, a different pattern was observed for NSP7- and NSP13-specific T cells. These cells were detected in only 3 out of 24 COVID-19 and in 2 out of 23 SARS recovered tested, but were present in 9 out of 18 unexposed donors (**Figure 4A/B**). The cumulative proportion of all studied subjects responding to NP and ORF-1-coded NSP7 and 13 proteins is shown in **Figure 4B**. These SARS-CoV-2 cross-reactive T cells from SARS-CoV-1/2 unexposed donors have the capacity to expand upon stimulation with SARS-CoV-2 peptides (**Figure 4C**). To better characterize the SARS-CoV-2 specific T cell reactivity detected in the SARS-CoV-1/2 unexposed individuals, fine-specificity and phenotype of the responding T cells were defined in selected donors. Characterization of the NP-specific T cells detected at high frequency in one donor (H-2) identified CD4 T cells reactive for an epitope comprised within the NP region 101-20. This same epitope was also detected in COVID-19 and SARS-recovered patients (**Figure 2B** and^8,18^). It has a high degree of homology to the MERS-CoV, OC43 and HKU1 NP sequences (**Figure 4D**). In two other SARS-CoV-1/2 unexposed donors (H-7 and H-3), we identified CD4 T cells specific for the NSP7 region 26-40 (SKLWAQCVQLHNDIL), and CD8 T cells specific for an epitope comprised within the NSP7 region 37-49 (NDILLAKDTTEAF), respectively (**Figure 4D, Extended Data Figure 3**).

**Fig 4:**
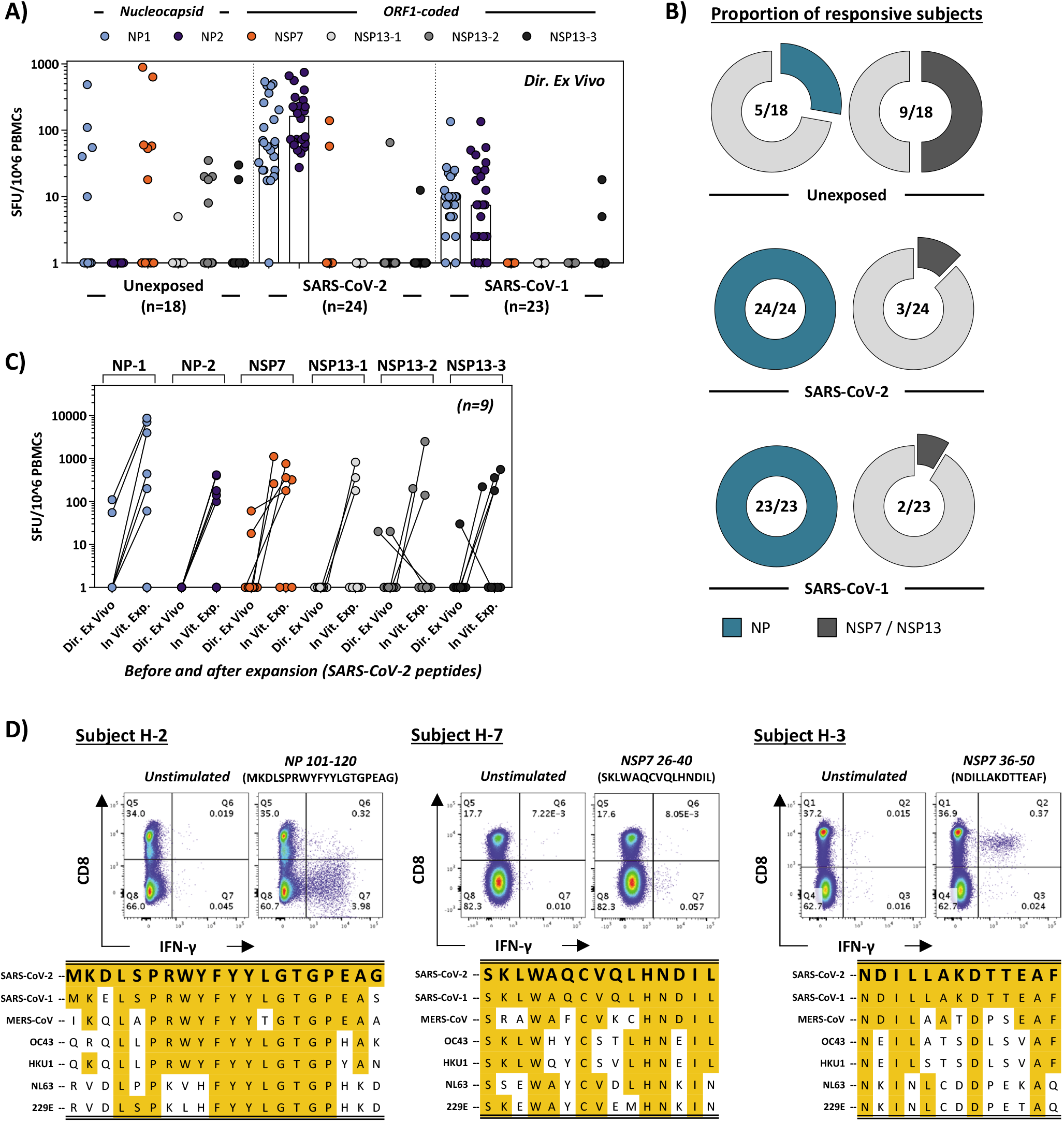
Differential protein immunodominance of SARS-CoV-2 specific T cells in COVID-19- and SARS-recovered patients and in unexposed individuals. (A) PBMC of individuals who are SARS-CoV-1/2 unexposed (n=18), recovered from SARS (n=23) or COVID-19 (n=24) were stimulated with peptide pools covering SARS-CoV-2 NP (NP-1, NP-2), NSP7 and NSP13 (NSP13-1, NSP13-2, NSP13-3) and analyzed by ELISpot. Frequency of peptide-reactive T cells is shown for each donor (dots) and the bars represent median frequency. (B) Pie charts represent percentage of individuals with NP-specific and NSP7/13-specific T cells for unexposed, SARS- and COVID-19-recovered individuals. (C) Frequency of SARS-CoV-2 reactive T cells in 9 unexposed donors to the indicated peptide pools directly ex vivo and after a 10-day expansion. (D) A peptide pool matrix strategy was applied in 3 SARS-CoV-1/2 unexposed individuals. The identified T cell epitopes were confirmed by ICS, and the sequences are aligned with the corresponding sequence of all coronaviruses known to infect humans.

These latter two T cell specificities were particularly intriguing since the homology between the two protein regions of SARS-CoV-1/2 and other “common cold” coronaviruses (OC43, HKU1 NL63 and 229E) was minimal (**Figure 4D**), especially for the CD8 peptide epitope. This may suggest that perhaps not only human “common cold” coronaviruses, but other presently unknown coronaviruses, possibly of animal origin, can induce cross-reactive SARS-CoV-2 memory T cells in the general population.

It was remarkable to find that NSP7/13-specific T cells were detected in 9 out of 18 (50%) SARS-CoV-1/2 unexposed donors, despite the fact that our analysis was performed with peptides that cover only 10% (684aa) of the ORF-1 proteome (7096aa). Notably, T cells specific for ORF-1-coded proteins were rarely detected in our SARS and COVID-19 convalescents. This is consistent with the findings of Grifoni et al^11^: using selected peptides, they detected ORF-1 specific T preferentially in some SARS-CoV-2 unexposed donors while T cells of COVID-19 recovered donors preferentially recognized structural proteins. The cause of this observed different pattern of immunodominance is presently unknown. We might speculate that a robust T cell response against structural proteins is induced by a productive infection (occurring in COVID-19 and SARS recovered patients). Individuals exposed to but not infected with possible unknown coronaviruses might just prime ORF-1-specific T cells. Indeed, induction of virus-specific T cells in “exposed but not infected” individuals has been demonstrated in other viral infections^20^. In coronavirus infected cells, the ORF-1 coded proteins are necessary for the formation of the viral replicase-transcriptase complex in which viral replication and transcription occur^14^. Therefore, an ORF-1-specific T cell can be envisioned to abort viral production in infected cells by lyses of SARS-CoV-2 infected cells even before the formation of mature virions.

Importantly, the ORF-1 region contains domains that are extremely conserved among many different coronaviruses^6^. The distribution of these viruses in different animal species might result in periodic human contact and subsequently induction of ORF-1-specific T cells with cross-reactive ability against SARS-CoV-2. Understanding the distribution, frequency and protective ability of the pre-existing structural or non-structural SARS-CoV-2 crossreactive T cells could be of great importance to explain some of the differences in infection rate or pathology observed during this pandemic. T cells specific for viral structural proteins have protective ability in animal models of airway system infection^21^,^22^. Nevertheless, the impact that the presence of ORF-1 specific T cells could have in the differential modulation of SARS-CoV-2 infection will have to be carefully evaluated.

## Material and Methods

### Ethics statement

All donors provided written consent. The study was conducted in accordance with the Declaration of Helsinki and approved by the NUS institutional review board (H-20-006) and the SingHealth Centralised Institutional Review Board (reference CIRB/F/2018/2387).

### Human samples

Donors were recruited based on their clinical history of SARS-CoV-1 or SARS-CoV-2 infection. Blood samples of recovered COVID-19 patients (n=24) were obtained 2 – 28 days post PCR negativity; of recovered SARS patients (n=23) 17 years post infection. Healthy donors’ samples were either collected before June 2019 for studies of T cell function in viral diseases (n=10) or in March-April 2020 and tested negative for RBD neutralizing antibodies and negative in an ELISA for NP IgG^19^.

### PBMC isolation

Peripheral blood mononuclear cells (PBMC) were isolated by density-gradient centrifugation using Ficoll-Paque. Isolated PBMC were either studied directly or cryopreserved and stored in liquid nitrogen until used in the assays.

### Peptide pools

15-mer peptides overlapping by 10 amino acids spanning the entire protein sequence of SARS-CoV-2 NP, NSP7 and NSP 13, as well as SARS-CoV-1 NP were synthesized (GL Biochem Shanghai Ltd; see **Sup. Tables 1,2**). To stimulate PBMC, the peptides we divided into 5 pools of about 40 peptides covering NP (NP-1, NP-2) and NSP13 (NSP13-1, NSP13-2, NSP13-3) and one pool of 15 peptides covering NSP7. For single peptide identification, peptides were organized in a matrix of 12 numeric and 7 alphabetic pools for NP, and 4 numeric and 4 alphabetic pools for NSP7.

### ELISpot assay

ELISpot plates (Millipore) were coated with human IFN-γ antibody (1-D1K, MabTech) overnight at 4°C. 400,000 PBMC were seeded per well and stimulated with pools of SARS-CoV-1/2 peptides (2 μg/ml). For stimulation with peptide matrix pools or single peptides, a concentration of 5 μg/ml was used. Subsequently, the plates were developed with human biotinylated IFN-γ detection antibody (7-B6-1, MabTech), followed by incubation with Streptavidin-AP (MabTech) and KPL BCIP/NBT Phosphatase Substrate (SeraCare).

### Flow Cytometry

PBMC or Expanded T cell lines were stimulated for 5h at 37°C with or without SARS-CoV-1/2 peptide pools (2 μg/ml) in the presence of 10 μg/ml brefeldin A (Sigma-Aldrich). Cells were stained with the yellow LIVE/DEAD fixable dead cell stain kit (Invitrogen) and anti-CD3 (clone SK7), anti-CD4 (clone SK3), and anti-CD8 (clone SK1) antibodies. Cells were subsequently fixed and permeabilized using the Cytofix/Cytoperm kit (BD Biosciences-Pharmingen) and stained with anti-IFN-γ (clone 25723, R&D Systems) and anti-TNF-α (clone MAb11) antibodies and analyzed on a BD-LSR 11 FACS Scan. Data were analyzed by FlowJo (Tree Star Inc.). Antibodies were purchased from BD Biosciences-Pharmingen unless otherwise stated.

### Cell culture for T cell expansion

T cell lines were generated as follows: 20% of PBMCs were pulsed with 10 μg/ml of the overlapping SARS-CoV-2 peptides for 1 hour at 37°C, subsequently washed, and cocultured with the remaining cells in AIM-V medium (Gibco; Thermo Fisher Scientific) supplemented with 2% AB human serum (Gibco; Thermo Fisher Scientific). T cell lines were cultured for 10 days in the presence of 20 U/ml of recombinant IL-2 (R&D Systems).

### Sequence alignment

Reference protein sequences for ORF1ab and Nucleocapsid Protein were downloaded from the NCBI database (see below). Sequences were aligned using the MUSCLE algorithm with default parameters and percentage identity was calculated in Geneious Prime 2020.1.2 (https://www.geneious.com). Alignment figures were made in Snapgene 5.1 (GSL Biotech).

| Accession ID | ORF1ab | Nucleocapsid Protein |
| --- | --- | --- |
| SARS-CoV-2 | QHD43415.1 | YP_009724397.2 |
| SARS-CoV-1 | NP_828849.2 | AAP33707.1 |
| MERS | YP_009047202.1 | YP_009047211.1 |
| OC43 | YP_009555238.1 | YP_009555245.1 |
| HKU1 | YP_173236.1 | YP_173242.1 |
| NL63 | YP_003766.2 | YP_003771.1 |
| 229E | NP_073549.1 | NP_073556.1 |

## Acknowledgments

We thank Mala K Maini (University College London, UK) and Subhash Vasudevan (EID, Duke-NUS Medical School) for critical reading of the manuscript. Grant support: Special NUHS COVID-19 Seed Grant Call, Project NUHSRO/2020/052/RO5+5/NUHS-COVID/6 (WBS R-571-000-077-733).

## Author contributions

NLB and ATT designed and performed experiments, analysed data, prepared the figures and edited the paper; KK, CYLT, MH, AC, ML, NT performed experiments and analysed data; MC, ML performed viral sequence homology and analysed data; WNC, LW provide antibody testing, MICC, EEO, SK, PAT, JGHL, YJT recruited patients and analysed data, YJT provided funding and AB designed and coordinated the study, provided funding, analysed the data, and wrote the paper.

---

**Extended Data Table 1:** Donor Characteristics

|  | **COVID-19**<br>**recovered** | **SARS**<br>**recovered** | **SARS-CoV-1/2**<br>**unexposed** |
| --- | --- | --- | --- |
| Number | 24 | 23 | 18 |
| Median age in years<br>(range) | 49.5<br>(27-78) | 49<br>(21-67) | 40<br>(33-63) |
| **<u>Gender</u>** |  |  |  |
| Male | 58% (14/24) | 26% (6/23) | 50% (9/18) |
| Female | 42% (10/24) | 74% (17/23) | 50% (9/18) |
| **<u>Residence</u>** |  |  |  |
| Singapore | 100% | 100% | 100% |
| **<u>Ethnicity</u>** |  |  |  |
| Chinese | 45.8% (11/24) | 43.5% (10/23) | 55.6% (10/18) |
| Caucasian | 37.5% (9/24) | 0% (0/23) | 27.8% (5/18) |
| Indian | 12.5% (3/24) | 21.7% (5/23) | 5.6% (1/18) |
| Japanese | 4.2% (1/24) | 0% (0/23) | 0% (0/18) |
| Malay | 0% (0/24) | 30.4% (7/23) | 11.1% (2/18) |
| Ceylonese | 0% (0/24) | 4.3% (1/23) | 0% (0/18) |
| **<u>\*Disease Severity</u>** |  |  |  |
| Mild | 66.7% (16/24) | 73.9% (17/23) | N/A |
| Moderate | 16.7% (4/24) | 13% (3/23) | N/A |
| Severe | 16.7% (4/24) | 13% (3/23) | N/A |
| Critical | 0% (0/24) | 0 | N/A |
| **<u>Virological parameters</u>** |  |  |  |
| SARS-CoV-1 PCR positive | N/A | 100% | N/A |
| SARS-CoV-2 PCR positivity | 100% | N/A | N/A |
| #SARS-CoV-2 NP Ig positivity | 100% | 100% | 0% |
| #SARS-CoV-2 RBD Ig positivity | 100% | 0% | 0% |
| Time since PCR negativity | 2-28 days | 17 years | N/A |

**\* WHO criteria**
**# reference: Yong et al., Lancet Infect Dis 2020**

**Extended Data Fig. 1:**
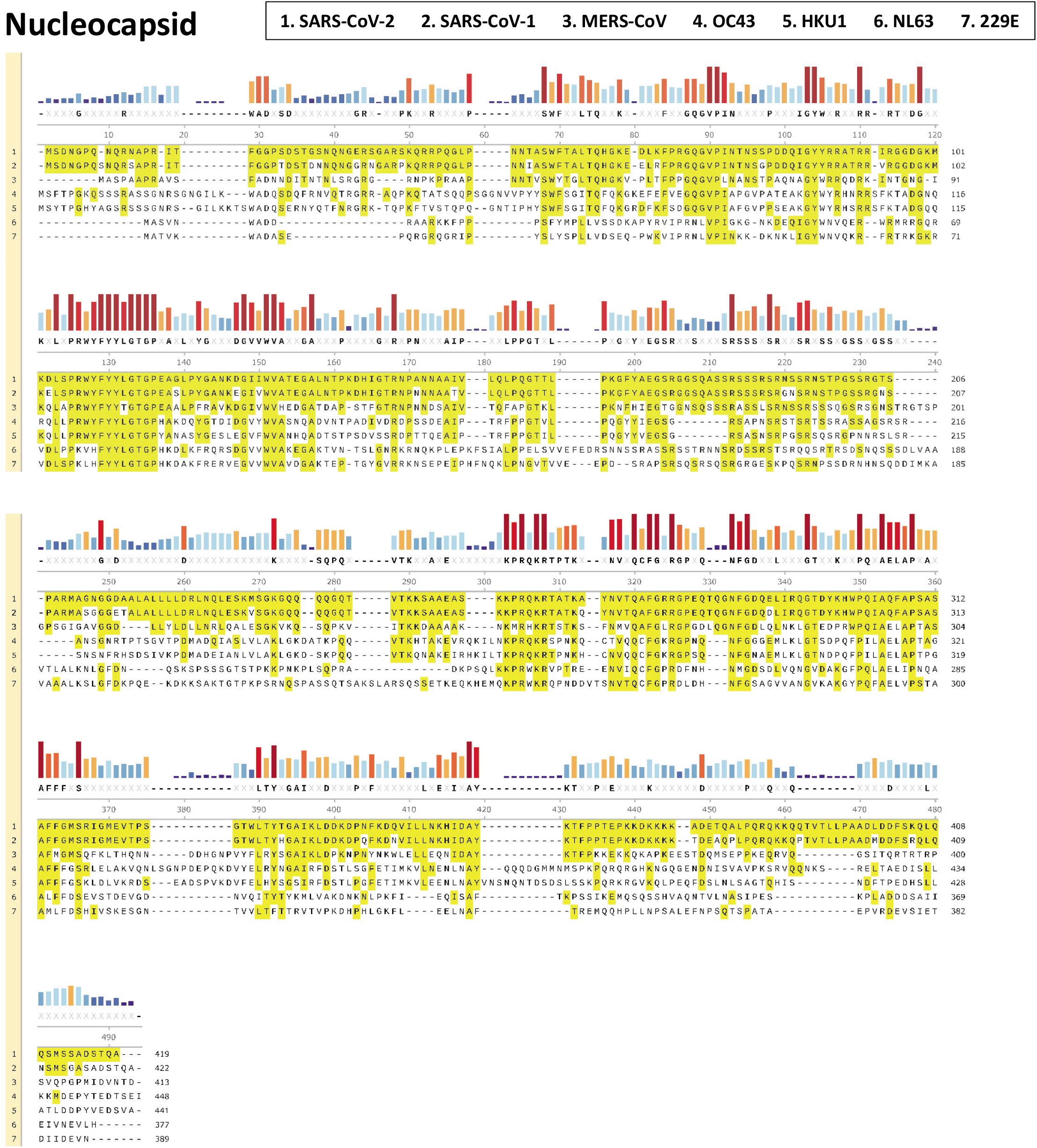
Sequence alignment of the nucleocapsid protein from all types of human coronaviruses. Amino acid sequences for Nucleocapsid Protein were downloaded from the NCBI database and aligned using the MUSCLE algorithm.

**Extended Data Fig. 2:**
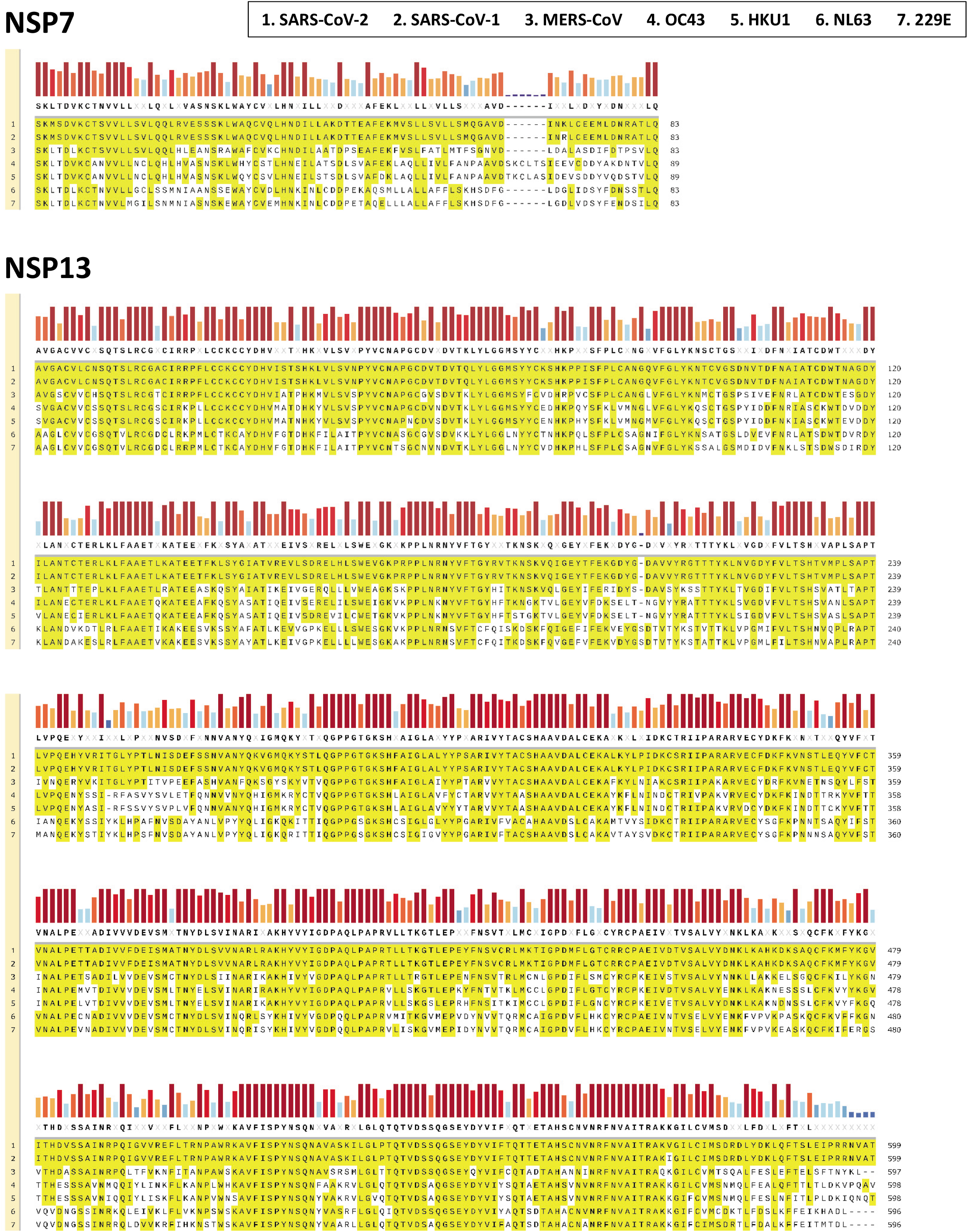
Sequence alignment of the ORF-1-coded nonstructural proteins NSP7 and NSP13 from all types of human coronaviruses. Protein sequences for ORF1ab were downloaded from the NCBI database and aligned using the MUSCLE algorithm. The alignment for NSP7 and NSP13 is shown.

**Extended Data Fig. 3:**
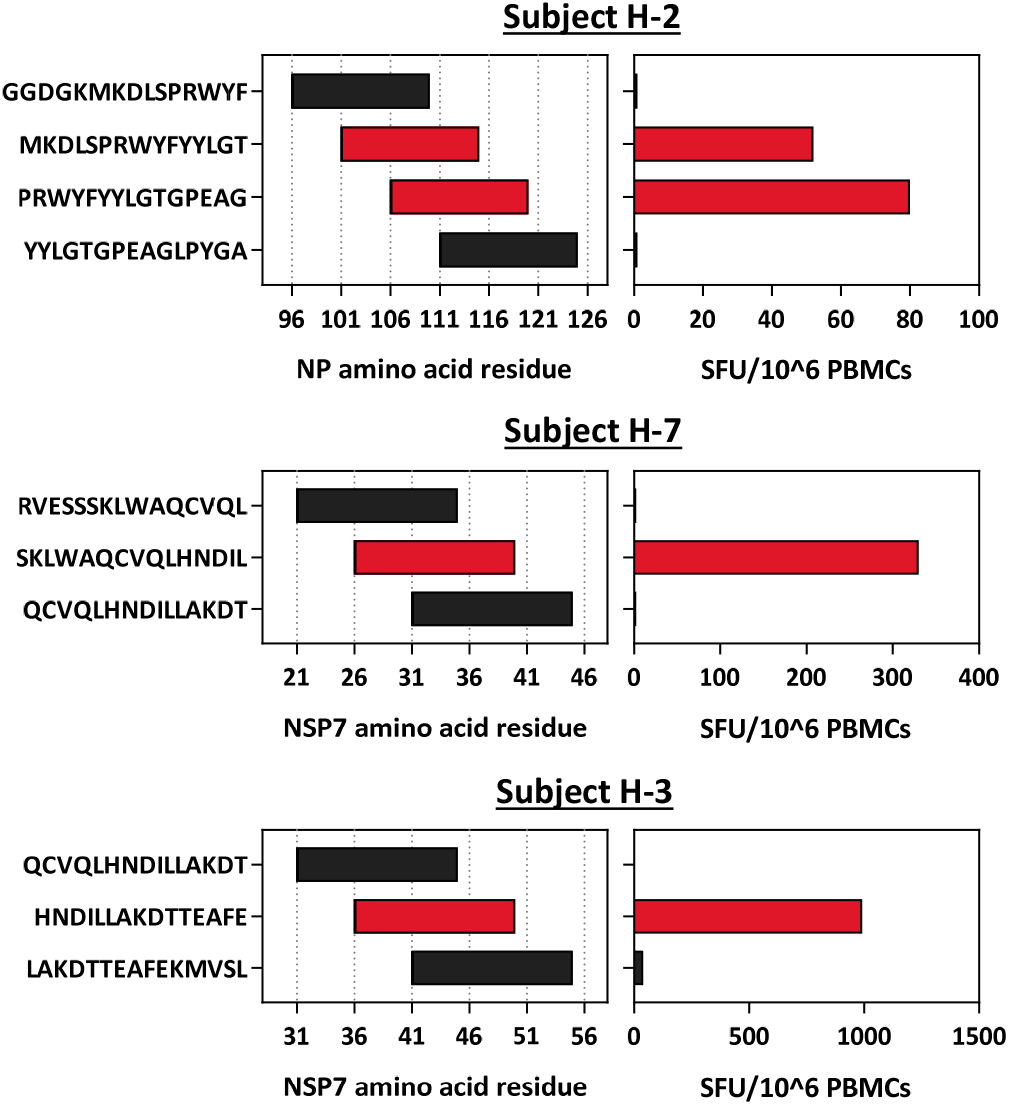
SARS-CoV-2 cross-reactive T cell epitope identification in three SARS-CoV-1/2 unexposed donors. PBMC were stimulated with the single peptides identified by the peptide matrix in parallel with the neighboring peptides and assayed by IFN-γ ELISpot. The amino acid residues are shown on the left; the frequency of IFN-γ-SFU/1 Million PBMC on the right. T cell-activating peptides in red, neighboring in black.

**Supplementary Table 1:**
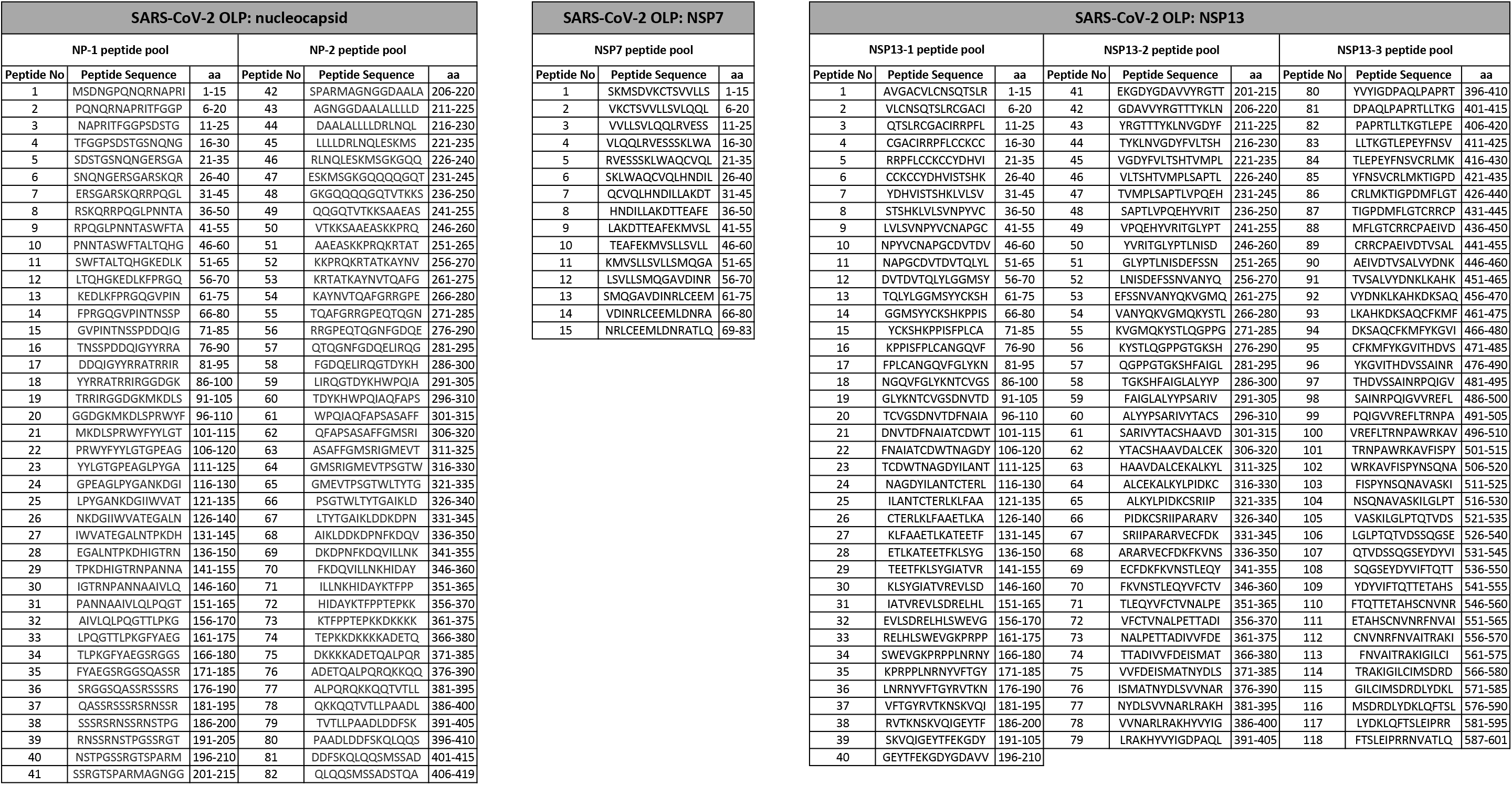
SARS-CoV-2 peptide libraries used in the study

**Supplementary Table 2:**
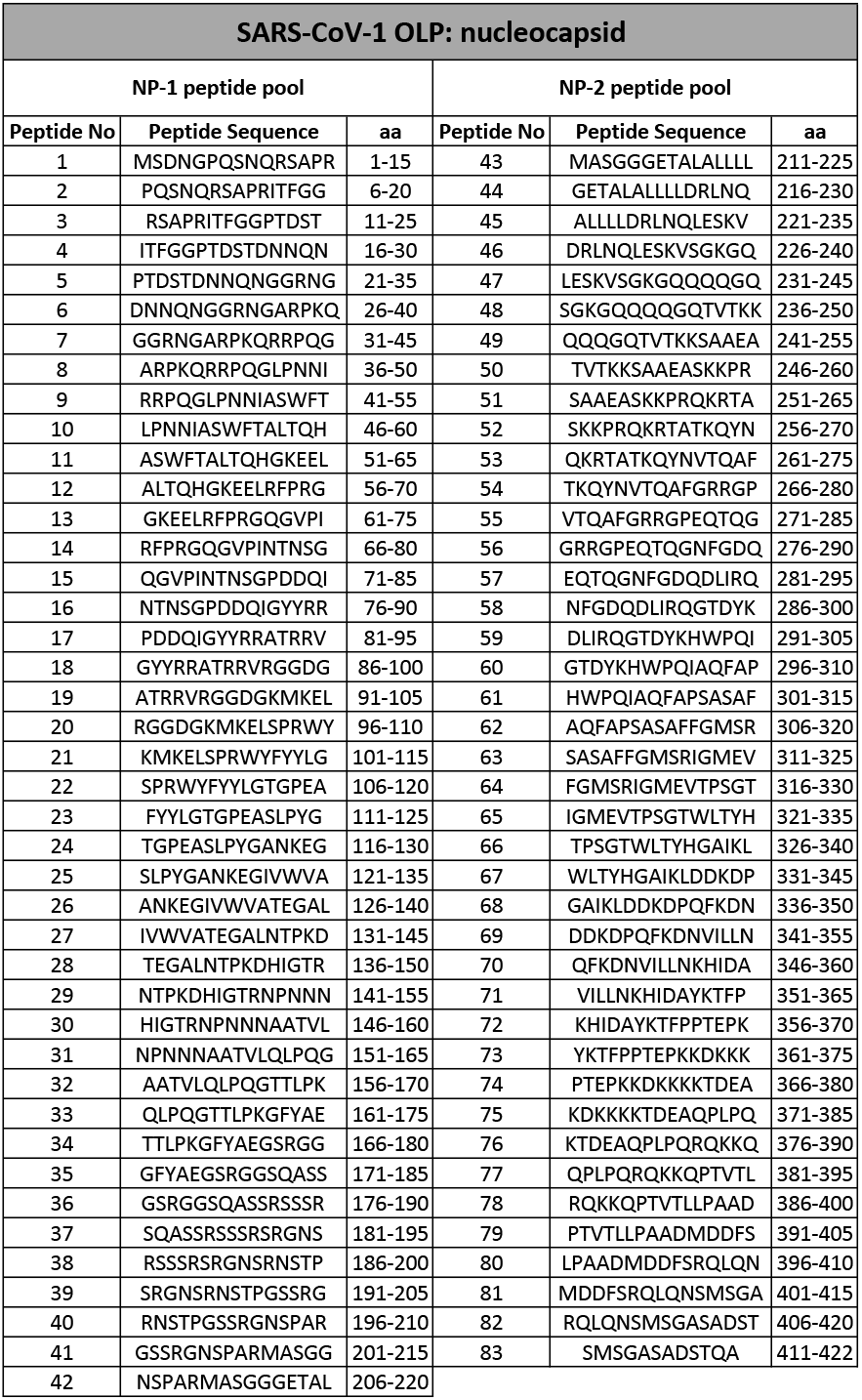
SARS-CoV-1 peptide libraries used in the study

